# The effect of boundaries on grid cell patterns

**DOI:** 10.1101/2020.05.16.099168

**Authors:** Mauro M. Monsalve-Mercado, Christian Leibold

## Abstract

Mammalian grid cells represent spatial locations in the brain via triangular firing patterns that tessellate the environment. They are regarded as the biological substrate for path integration thereby generating an efficient code for space. However, grid cell patterns are strongly influenced by environmental manipulations, in particular exhibiting local geometrical deformations and defects tied to the shape of the recording enclosure, challenging the view that grid cells constitute a universal code for space. We show that the observed responses to environmental manipulations arise as a natural result under the general framework of feedforward models with spatially unstructured feedback inhibition, which puts the development of triangular patterns in the context of a Turing pattern formation process over physical space. The model produces coherent neuronal populations with equal grid spacing, field size, and orientation.

**PACS numbers:** 87.19.lv,87.10.Ed,02.30.Jr

## I. INTRODUCTION

Grid cells in the medial entorhinal cortex (mEC) of rodents fire whenever the animal walks over the vertices of an idealized triangular lattice [1]. As a population, the grid cell network is thought to provide an efficient neuronal representation of spatial locations in the environment [2, 3], and moreover, it is a prime candidate for the biological implementation of path integration [4, 5] and path planning [3]. However, many experimental reports argue against grid cells representing a perfect triangular lattice. They exhibit field-to-field amplitude and location variability, lattice defects, geometrical distortions of the triangular pattern, and respond strongly to environmental manipulations and the geometry of the recording enclosure [6–14]. These observations challenge the role of grid cells in path integration and their pivotal role in encoding of spatial information.

While there is only limited and indirect experimental evidence for the involvement of grid cells in path integration [15, 16], the currently overwhelming support for this idea arises from mechanistic computational models, which show that integrated velocity information can be stored in the grid cell population, either by means of continuous attractor states [17] or by a superposition of velocity controlled oscillatory inputs [5].

More recently, however, a third class of mechanistic models has come into focus that does not necessarily link grid patterns to path integration and thus opens an alternative view on the functional role of grid cells. These models explain the development of grid patterns by assuming space selective inputs that undergo experience-dependent synaptic plasticity. A variety of these *feedforward* models have been proposed that are all based on different biological assumptions, such as neuronal adaptation [18, 19], excitatory-inhibitory balance during synaptic plasticity [20, 21], non-linear contributions to synaptic learning rules and different ratios of potentiation and depression [22], and a combination of hippocampal phase precession and spike-timing dependent synaptic plasticity (STDP) [23].

Despite these divergent mechanistic explanations, feed-forward models explain the development of hexagonal firing patterns by the same mathematical principles of a Turing pattern formation process [24, 25]. Following this view, we employ a simple pattern formation formalism to capture the essence of feedforward models. In this work, we study the role of static and spatially unstructured feedback inhibition on the pattern formation process. We show that feedforward models with additional unspecific feedback inhibition can explain a number of experimental observations. First, given enough time, grid cells can robustly learn perfect triangular patterns in recording enclosures of arbitrary shape without assuming special treatment for the boundaries. Moreover, the grid cell network exhibits coherent responses such as identical spacing, field size, and orientation, without underlying local attractor dynamics.

In contrast to experimental reports of homogeneous distributions of grid phases [26], our model produces grids belonging to three phase clusters, which we argue can be resolved by prematurely terminating synaptic learning. Phase clustering in our model, however, predicts that in future experiments with many spatially resolved grid cells, the phases may not be strictly uniform, but show biases towards three preferred phases.

Finally, we show that most of the observed defects, distortions, and responses of the grid cell pattern to polarized enclosures and environmental manipulations, can also be understood as a result of unfinished learning. In addition, we reproduce some of these observations without special considerations beyond the shape of the recording enclosure.

These results suggest that feedforward models with inhibitory feedback are a natural framework to study the development of triangular patterns in grid cells.

## II. RESULTS

We study the Turing-type pattern formation process implemented by feedforward models with a simple formalism still capable of explaining the influence of boundaries (Fig. 1A). Activity *E* of entorhinal cortex cells is assumed to result from rich spatial information coming from a population of spatially modulated cells with activities *H* that are typically, but not necessarily, assumed to be hippocampal place cells. In a firing rate approximation, the contribution from activities *H* are weighted by effective connections *W* that determine the dynamical evolution

**FIG. 1.**
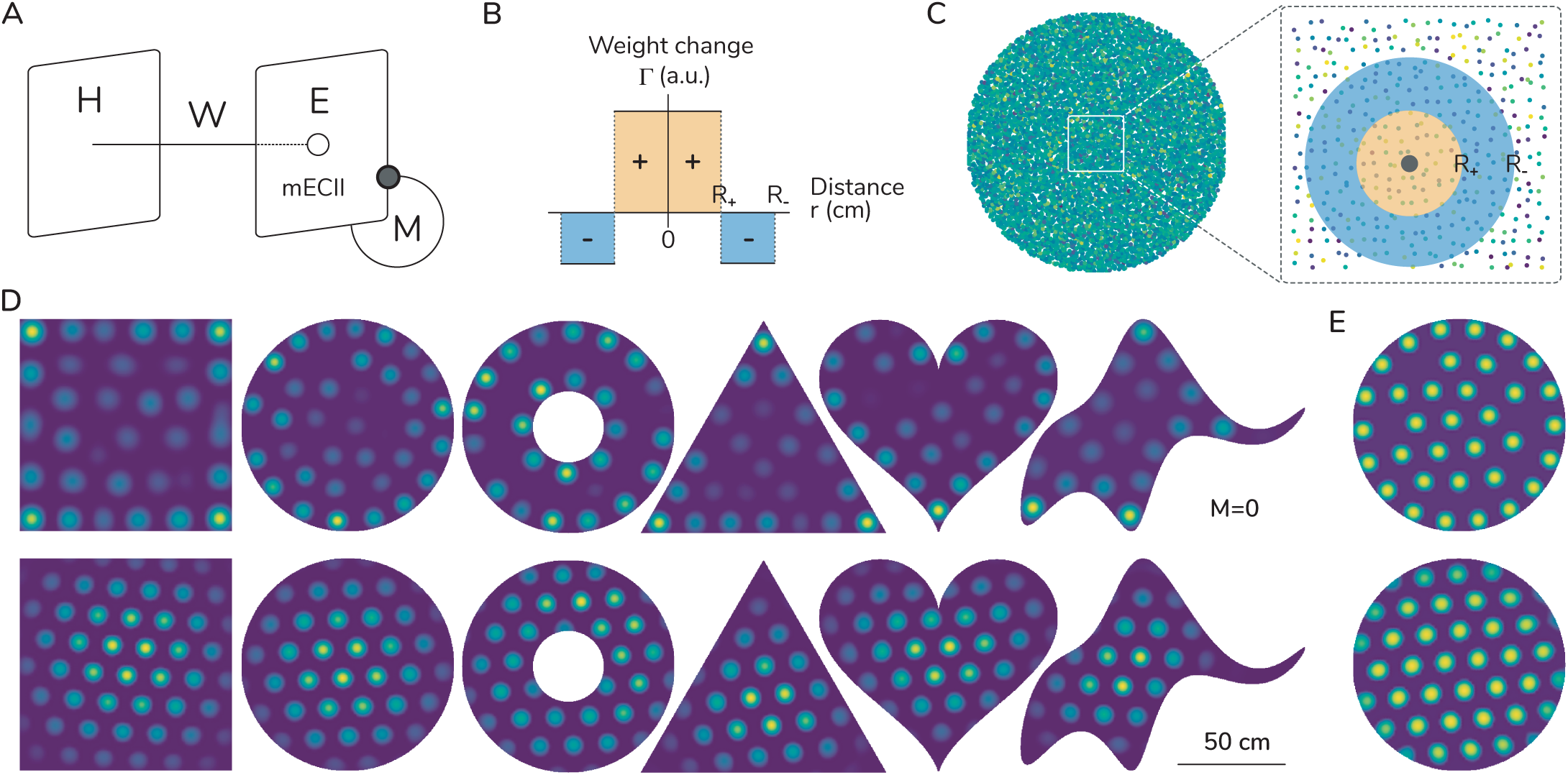
Feedback inhibition allows the development of triangular patterns inside enclosures of arbitrary shape. (A) A layer of cells with spatially selective activity *H* projects with effective weights *W* to cells *E* presumably in the entorhinal cortex. In this work, entorhinal cells are connected via a random inhibitory coupling *M* with no spatial structure or dependence on the boundaries. (B) Hebbian learning is implemented by an update rule for the weights that depends on their spatial distance. The effective learning kernel Γ is always of the center-surround type, here captured by a general step function of distance. (C) The learning rule implements a Turing pattern formation process on the development of the weights *W* (*x*) (greenish dots). A weight in a certain location (black dot) increases in proportion to weights in a neighbouring area (yellow), while it decreases in proportion to weights in a further surrounding region (blue). (D) Simulations of the growth process for weights (green) constrained to enclosures of different shape reveal that when no feedback inhibition is present (top row), the weights tend to cluster along the boundaries with a preference for corners and regions of high curvature. On the other hand, the presence of feedback inhibition (bottom) produces triangular patterns in all shapes without special treatment of the boundaries, i.e. without the use of periodic or fading boundary conditions. (E) Pure Hebbian learning is inherently unstable with weights growing without bounds. An additional stabilizing non-linearity only used in this panel after 600 time steps of learning reveals that, far from the boundaries, weights may still form local triangular patterns, even when no feedback inhibition is present (top). With feedback inhibition (*M ≠* 0) the additional non-linearity removes the apparent fading close to the boundaries observed in panel D.

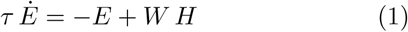

of entorhinal cells’ activity on a fast neuronal time scale *τ*. The effective weights *W* experience plastic changes according to a general linear Hebbian learning rule

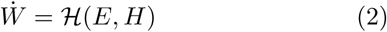

depending on the activity of pre- and postsynaptic cells that evolves in a slow time scale compared to the cellular dynamics (*τ «* 1). We thus can separate time scales and use the equilibrium rates *E* = *WH* for computing the weight updates [27]. Importantly, and consistent with biology, weights are restricted to remain positive, ensuring that patterns tend towards hexagonality. As examples for ℋ, a pure Hebb rule ℋ= ⟨ *E*^*T*^ *H* ⟩ would be implemented by time-averaged cell co-activation, while an STDP rule ℋ = ∫ d*s A*(*s*) ⟨ *E*^*T*^ (*s*)*H*(*t* −*s*) ⟩ weights causal relationships by a learning window *A*(*s*) [27–29]. The operator ℋ could also include non-linear terms as they, for example, would arise from a BCM rule [30]. Each of the different feedforward models involves a specific choice of learning rule and additional biological mechanisms, that transform the learning equation into a Turing-type pattern formation process [25, 31] on the synaptic weights, where the pattern-forming contribution can be described as

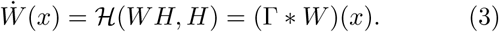

Additional non-linear local terms in *W* are typically included to stabilize the growth process, and are often only relevant at late stages in the development of the patterns [32].

The Turing-type learning rule describes the growth process of weights associated to a location *x* inside a bounded enclosure of arbitrary shape. The growth of weights is dictated by their convolution with the learning kernel Γ, whose shape follows a general center-surround profile (Fig. 1B), resembling those used by the various feedforward models in the literature [19, 21–23]. To motivate this learning rule we can think of each location *x* inside the recording enclosure as being encoded by the corresponding input activity *H*(*x*) and subsequently transferred to entorhinal cells by the corresponding weight *W* (*x*). This weight grows proportionally to the summed contribution of weights *W* (*y*) located in a nearby region (*| y* −*x | < R*_+_), whereas it decreases in proportion to weights *W* (*z*) in a further-surrounding area (*R*_+_ *< |z* −*x |< R*_−_, Fig. 1C). How much the neighboring weights contribute to the change of *W* (*x*) is encoded in the shape of the learning kernel Γ(*x*).

Numerical simulations of the growth process dictated by Eq. 3 reveal a tendency for weights to develop into spatially defined fields of high strength (Fig. 1D, top row). For enclosures with boundaries of arbitrary shape, and without the use of artificial boundary conditions such as periodic or fading boundaries, there is a general trend for weights to cluster and be stronger along the boundary, emphasizing certain locations where firing fields will be strongest depending on the geometry of the enclosure. In particular, corners and regions of high concavity promote the development of strong weights.

Clustering of fields nearby the physical boundaries of a Turing system is an intrinsic feature of the pattern formation process. It can be observed in nature whenever a physical obstruction to the rearrangement of activity is present, for instance in experiments involving chemical reaction-diffusion systems in a dish [33]. Grid cells are unusual in this respect because experiments do not report such strong linkage to boundaries and thus pure feedforward Turing-type pattern formation seems not to be able to explain grid cells in bounded enclosures.

### A. Feedback inhibition allows the development of triangular patterns inside polarized enclosures

We now introduce additional feedback inhibition to the model and explore its effects on the learning process. To this end, entorhinal cells are coupled among each other by means of an inhibitory connectivity matrix *M*, which modifies the firing rate dynamics to

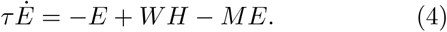

Importantly, the connectivity matrix *M* is not subject to any plastic changes and remains unaltered during the learning process. Its entries are randomly drawn from an arbitrary positive distribution such that each cell receives feedback inputs with mean total synaptic strength *m*_0_ and variance *σ*^2^ (Methods). In particular, the choice of *M* does not introduce any dependence on the spatial geometry of the enclosure or the spatial or functional arrangement of cells in the cortex.

Numerical simulations of the learning process reveal how the development of synaptic weights changes when grid cells are strongly coupled via inhibition (Fig. 1D, bottom row). In contrast to the uncoupled case, weight patterns always reach a perfect triangular-lattice arrangement of fields that cover the entire enclosure independent of the particular geometry of its boundary. The strength of the triangular patterns is weaker close to the boundary. It is a consequence of the inhibitory coupling in the learning process and does not result from fading boundary conditions nor any other special considerations for the boundaries like, e.g. border cell inputs [34, 35].

The results reported here do not depend on the specific stabilizing biological mechanism stopping the learning process. In particular, weights fade out at the borders because we did not include any additional prescription to stop their unbounded growth. For both the uncoupled and coupled cases, the fading around the center of the enclosure or towards the boundary disappears when a soft upper bound is activated after the triangular pattern has been learned (Fig. 1E).

### B. Boundaries introduce a growth bias

To analyze how feedback inhibition weakens boundary dependence, we first explore the effects of boundaries without inhibition. The simulated patterns presented in the top row of Fig. 1D, as well as experimental descriptions of Turing systems [33], point towards an inherent proclivity of the growth process to develop stronger activity nearby boundaries. The way in which the growth process affects different locations within the enclosure reveals the key role of boundaries. Weights located far away from the edge follow the description of Fig. 1C. They sense the added effect of weights located within the potentiating and depressing regions. In comparison, weights nearby the edge sense the effect of an incomplete ring of depression, thus being effectively more potentiated compared to weights far from the edge. Weights nearby the edge are thus missing the depressing effect of weights located outside the enclosure. In biological terms, during an open field exploration task, if an animal remains inside the recording environment there is no activity of place cells encoding locations outside. As a consequence, entorhinal cells are excited only by place cells inside the environment, thereby introducing the growth bias into the Hebbian weight dynamics.

Fig. 2A illustrates the restriction imposed by the boundary on the growth process, i.e., the spatial range of action of the learning rule is limited by the physical extent of the enclosure. The growth bias ß (Materials & Methods) measures the total contribution that weights in a surrounding region add to the growth of a weight at the center. Its impact is proportional to the overall effect of the truncated learning kernel, corresponding to its integral within the enclosure. Fig. 2B shows how its integral changes with respect to the distance from its center to an infinite straight boundary. Far away from the edge the learning kernel is complete and therefore its integral remains constant. Once it comes in contact with the edge, the progressive truncation of its depressing component results in an increasing effective potentiation. The illustrated principles are independent of the particular shape of the learning kernel, always introducing a higher bias near the edge.

**FIG. 2.**
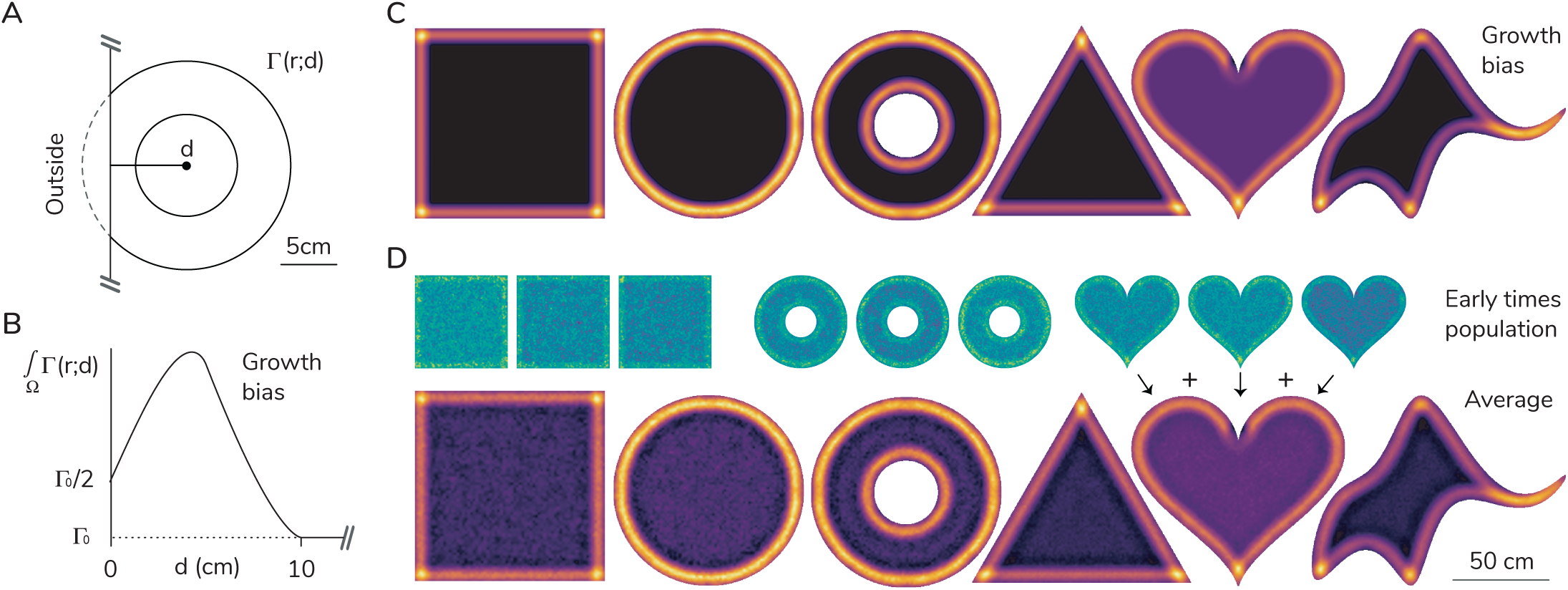
Physical boundaries introduce a bias into the growth process. (A) Weights close to the boundary are potentiated more compared to far away weights inside the enclosure. A weight (black dot) closer than the spatial range of action of the learning kernel loses a region of depression that would come from activity of cells located outside the enclosure. (B) The result is a bias ß (Methods) in the rate of growth of weights depending on their distance d away from the boundary. It is proportional to the overall effect (the integral) of the learning kernel centered at d and constrained to the inside of the enclosure. (C) The growth bias computed numerically for all shapes predicts the locations where weights are predisposed to grow the most. (D) Simulations of the growth process for the case when no inhibition is present reveals that at early times the individual patterns (green) reflect the predicted growth bias. Since cells are uncoupled they represent different random instances of the growth process. Their average (red) approximates the predicted growth bias, even for later stages of the process.

We numerically computed the growth bias for the enclosures depicted in Fig. 1D. They predict a location specific bias depending on the particular geometry of the boundary (Fig. 2C). The growth bias agrees with the results obtained for the patterns in Fig. 1D, leading to higher activity near corners and concave regions. In addition, activity splits into discrete fields of activity. The specific location of these fields depends in general on the initial configuration of the system, however, regions with growth bias excess will attract over-proportionally more firing fields. To show that the prediction from Fig. 2C matches the real growth bias, we simulated early-time weight development of a population of grid cells when no inhibition is present (Fig. 2D, green), and computed the corresponding population averages (Fig. 2D, red). Because the cells are uncoupled, each pattern represents a different realization of the growth process for different initial random states of the system. As a consequence, their population average reflects the underlying growth bias which excellently agrees with the predictions shown in Fig. 2C.

### C. Inhibition produces coherent populations

The previous analysis reveals that an uncoupled population of entorhinal cells is able to encode the underlying growth bias in a robust way. This is suggestive for the mechanism implemented by strong inhibitory coupling to remove the growth bias from the pattern formation process. Indeed, in the presence of homogeneous inhibition (all cells inhibit each other with exactly the same strength) the Turing learning prescription (Eq. 3) for each entorhinal cell is modified to

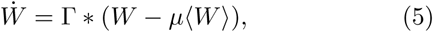

where ⟨ *W* (*x*) ⟩ denotes the population average of weight patterns at position *x* (see Methods). The unbiasedness parameter *µ* := *m*_0_*/*(1 + *m*_0_) controls the relative effect of the average ⟨ *W* (*x*) ⟩ on the weight change. It depends strongly on the synaptic amplitude *m*_0_ of the inhibitory connectivity matrix; it vanishes when no inhibition is present, and asymptotically approaches 1 for increasing inhibitory strength. The unbiasedness reflects the ability of the inhibitory network to correct for the bias introduced by the boundary such that the pattern never fails to reach a triangular pattern for unbiasedness greater than zero, however convergence times are often much longer for weaker unbiasedness; see below.

### D. Time course of pattern formation

Subtracting the population average from the weight dynamics makes the overall magnitude of weights grow slower independent of the convergence time to reach triangularity, however, it does not stabilize the learning dynamics and weights will continue to grow unbounded at an exponential rate. It naturally allows patterns to develop before additional stabilization mechanisms come into effect. Despite this unbounded exponential growth, the coefficients of variation (CVs) along the spatial as well as the grid cell dimensions reach a saturation point early on in the pattern formation dynamics (Fig. 3A), pointing towards the growth of an extended spatial structure and no winner-take-all dynamics at the population level. This is a direct consequence of the non-negativity constraint on the weights, with CVs growing exponentially when no restrictions are imposed on the weights.

**FIG. 3.**
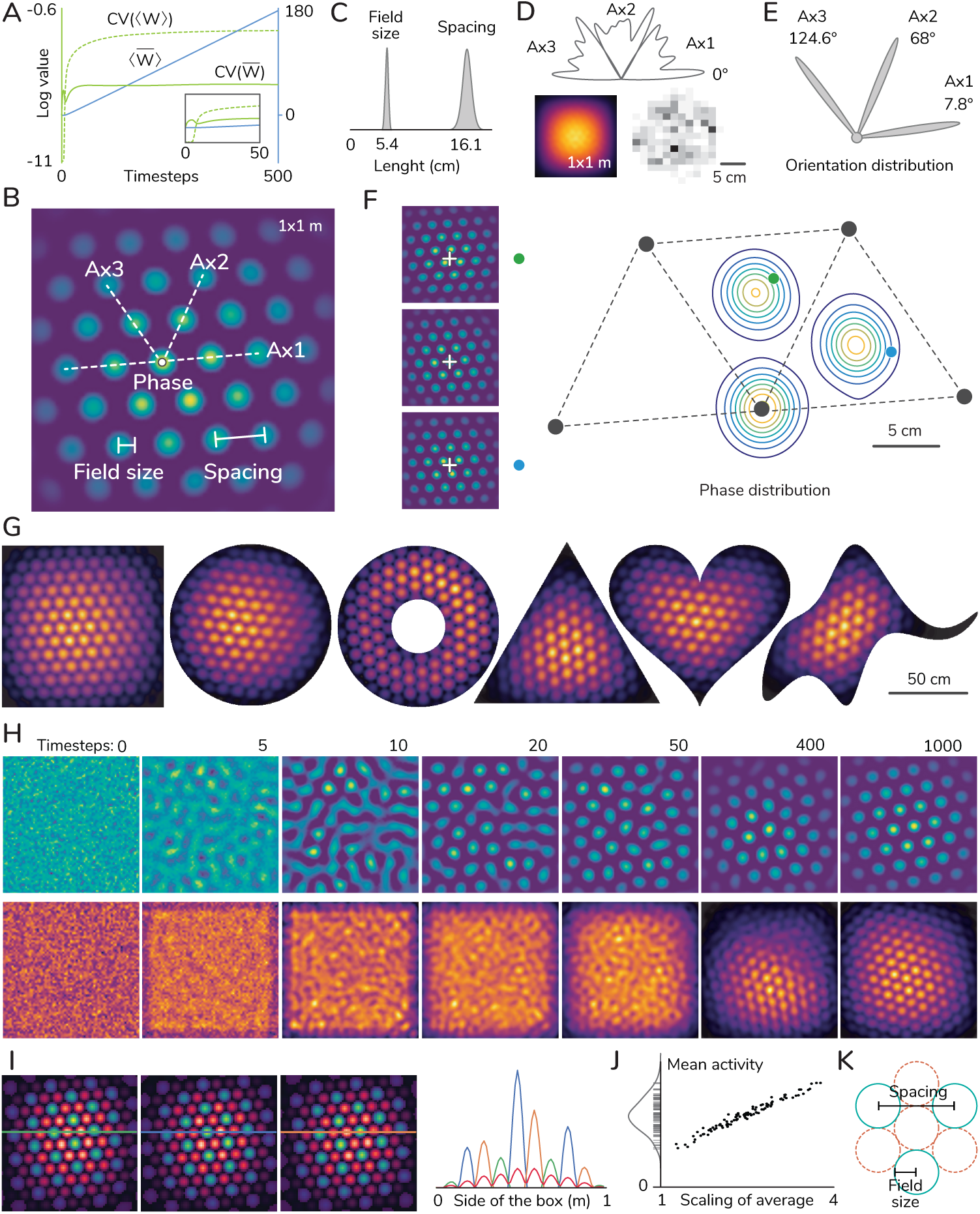
Feedback inhibition produces coherent population responses. (A) Simulations of the growth process for the coupled case (strong feedback inhibition) reveal that the average of patterns over space (bar) and over grid cells (angular brackets) grows exponentially in time (blue curve, semi-logarithmic scale). In contrast, the coefficient of variations (green curves) both of the spatial averages 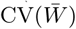 (i.e. along the grid cell dimension) and of the population average CV(⟨ *W* ⟩) (i.e. of the spatial distribution) saturate to a constant value early in the growth process (see inset). (B) Definitions of the local geometric measures used to characterize the grid cell pattern. (C) Population distribution of the cell averaged measures of field size and spacing. The ratio of field size to spacing is consistently approximately 1*/*3 for all parameters explored. (D) For a population with homogeneous connectivity, the orientations (top) and spatial phases (bottom) are uniformly represented. Bottom shows the population average (left, red) and a histogram of the field centers choosing only the fields closest to the midpoint of the square. (E) The distribution of orientations for noisy connectivity shows that all patterns share a preferred orientation. (F) For noisy connectivity, patterns self-organize into three evenly distributed phase groups (contour plot). The skeleton represents the nearest neighbouring fields (vertices, black dots) of a pattern belonging to one of the phase groups. The left column shows example patterns for each of the groups (as marked by the green and blue dots); the middle pattern is from a reference group (black). The white crosses mark the center of the enclosure. (G) Population averages (red) suggest that patterns self-organize into phase groups independent of the geometry of the enclosure. (H) The time series evolution of a single pattern (green) and the population average (red) illustrates the self-organization into phase groups and the development of a perfect triangular pattern. (I) Three patterns from different phase groups (green) on top of the population average (red) for weak connectivity strength (*µ* = 0.1, only in this panel) illustrate the approximate non-overlapping nature of the phase groups. Slices of activity of the three patterns (green, blue, and orange lines) show that the population average at any position is mostly governed by one phase group. (J) The patterns’ mean activity (vertical axis) is approximately normally distributed. For each pattern we computed the scaling factor by which the average activity needs to be multiplied to fit the non-zero regions of the pattern (in a minimal mean square sense; all fitting errors are below 1% of each pattern’s peak). (K) For packed tangent circles mimicking perfectly non-overlapping phase groups, the ratio of field size to spacing corresponds to 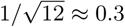.

Fig. 3B defines geometrical measures such as spacing, orientation, and spatial phase, that are used to characterize grid cell patterns [1]. We investigated the dependence of these measures on the inhibitory connectivity. The population of grid cells shares the same spacing independent of the strength of the inhibition (Fig. 3C and Mov. S1-5), since it only depends on the shape of the learning kernel Γ [36]. For perfect homogeneous connectivity the population represents all orientations and phases uniformly (Fig. 3D and Mov. S6). However, small perturbations of the connectivity send the system to a qualitatively different state: adding a small amount of randomness to the coupling *M* breaks the symmetry and, as experimentally found [1], produces a coherent population that shares the same orientation (Fig. 3E). The selected orientation generally depends on the geometry of the enclosure (see examples in Fig. 1D). A biologically reasonable way of inducing random deviations from uniform coupling is to consider random sparse connectivity of equal strength. With such a connectivity matrix we obtain qualitatively similar grid populations for sparseness values ranging from totally connected (close to 100%) to very low sparsity levels (3%) (see Mov. S7 for 10% sparseness).

Moreover, noise in the connectivity promotes random lateral spread that ensures the rapid correction of lattice defects and might help learning new representations faster. Conversely, patterns learned with homogeneous connectivity are more stable, which diminishes their ability to correct for defects present at early times (Mov. S6). Furthermore, the contribution of noise to the growth rate comes in the form of a random average of all patterns (Materials & Methods) which in general exhibits hexagonal symmetry even when patterns have a uniform distribution of phases and orientations. As a result the most spatially stable configuration (least lateral spread of the fields) is the one where fields are proportional to the population averaged activity.

The latter consideration explains a striking feature of the learned grid patterns, viz., that their spatial phases converge to clusters of three dominant phases (Fig. 3F). They correspond to the three phases needed for the patterns to evenly cover the entire environment while reducing the overlap among patterns from different phase groups. The organization of the patterns into three roughly non-overlapping phase groups is robust to noise in the connectivity, since the growth of a single pattern receives strong feedback from the population, and thus patterns can only reach periods of spatial stability together. The observation of three coherent phase groups is not limited to the square enclosure, they can be observed for all shapes investigated (Fig. 3G). Experimental reports do not support a clear clustering of phases into three groups [26]. This can, however, be accounted for by our model under the assumption that synaptic plasticity stops before three clear clusters have evolved (Figure. 3H, and next section).

The model in the simple form presented here (without stabilizing homeostatic mechanisms) does not produce asymptotically stable spatial patterns (see all supplementary movies). The population arrives at the three-phase organization depicted in Fig. 3F,G as patterns first reach triangularity (Fig. 3H). After a period of spatial stability, the population average undergoes transitions through disordered states before returning back to the three-phase organization. At any point in time, in the absence of a stabilizing non-linearity, individual cells’ grid patterns may undergo quick transitions from one phase group to another, followed by a period of spatial stability (see Mov. 1-5).

Since structure formation is strongly guided by the phase clusters, our model predicts that even if the clusters are not clearly visible in experiments, phases should at least not be uniform. Therefore we next explore functional consequences of the phase clusters. To this end, we visualized grid cell activity in relation to the population average (Fig. 3I; for small unbiasedness *µ* to obtain more concentrated phase distributions) to show that patterns from different clusters minimize their overlap while maintaining good coverage of the whole space. It follows naturally from the organization of the patterns into roughly non-overlapping groups that the ratio between field size and spacing must match the experimentally reported value of about one third, since, for three densely packed phase clu<u>ste</u>rs, the ratio can be geometrically deduced to be 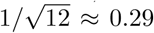 (Fig. 3K). Dense packing of phase clusters does not depend on small *µ*. In Fig. 3J, we show that, for large *µ* = 1, where fields get slightly broader, the population average at any position is still mostly determined by one phase cluster, as scaling factors between field activity and population average concentrate around 2 to 3, meaning that on average roughly a third of neurons (one phase cluster) are active at any position despite the broad distribution of the cells’ mean activities. Thus clustered phases seem to not strongly deteriorate homogeneous coverage of space by grid fields.

### E. Unfinished learning explains distorted patterns

Contrary to the patterns produced by most models of grid cells, experimentally obtained grid patterns hardly ever are perfectly triangular lattices. A number of experimental reports investigate typical deviations from a perfect pattern, including lattice defects, geometrical distortions, and particular responses to environmental manipulations [7–14]. In the realm of feedforward models with feedback inhibition, all of these effects can be the result of unfinished learning. Moreover, assuming such premature termination of synaptic plasticity, we also can account for homogeneous distribution of spatial phases (columns 4 or 5 in Fig. 3H) as well as stability of grid fields.

Distortions reflect different aspects of the Turing pattern formation process. We reproduce with our model some of these observations without introducing any additional details beyond choosing the geometry of the enclosure (Fig. 4, see details in the figure legend).

**FIG. 4.**
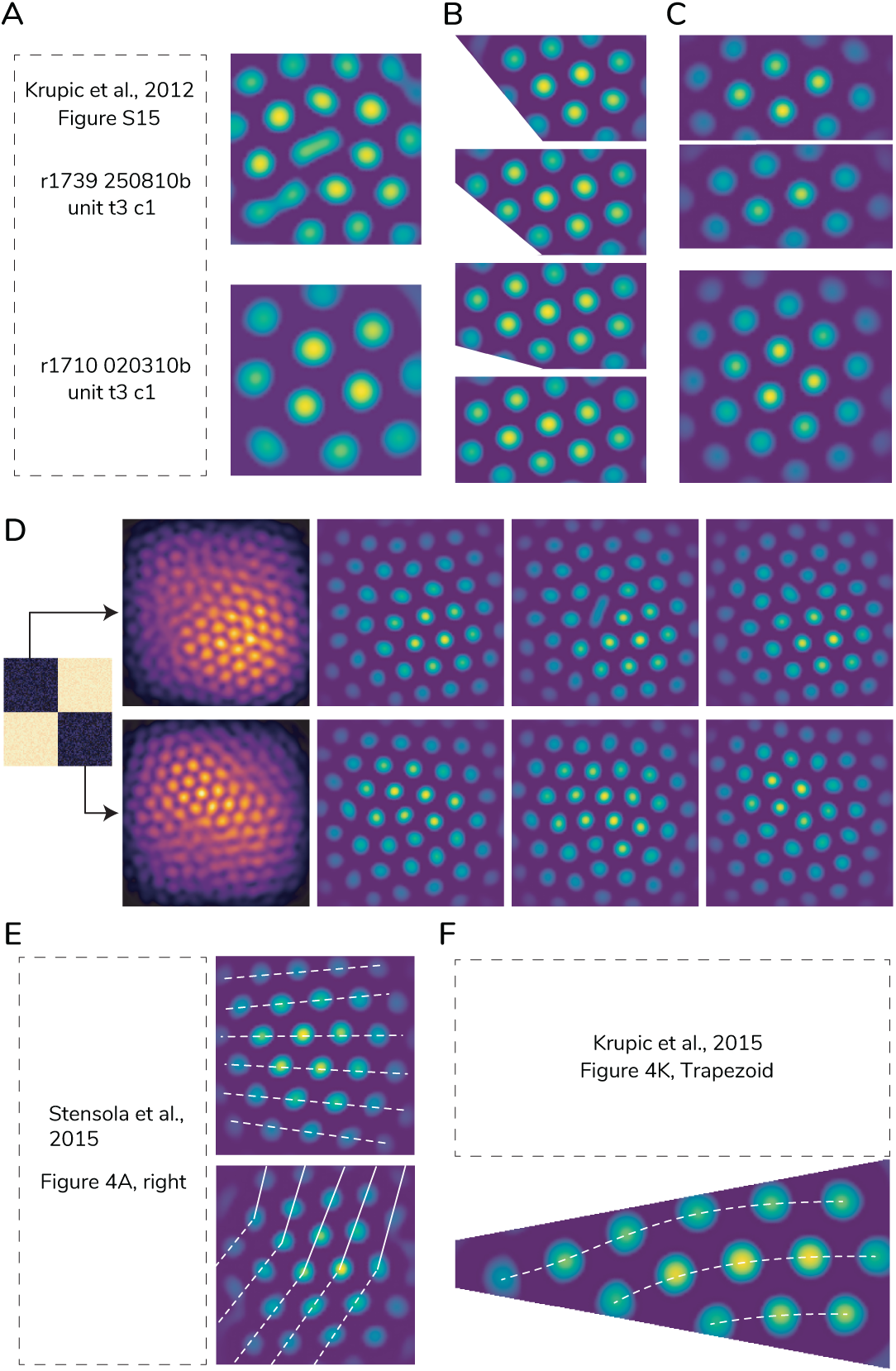
The effects of unfinished learning on the grid cell pattern. (A) Typical defects found during a Turing pattern formation process are splitting fields (top left) and lattice dislocations (penta-hepta defect; bottom left). Examples match experimentally obtained rate maps from grid cells [12]. We simulated the developments of patterns on a square enclosure from a random initial state. Stopping learning before spatial stability is achieved results in similar defects on some of the patterns (right column). (B) Extending the recording enclosure reveals the appearance of a consistent pattern [10, 13]. We simulated the pattern in an initial trapezoid until it was spatially stable. After expanding the trapezoid we continue learning the whole pattern, with the novel region starting from an initial random state. The new fields become a coherent part of the pattern learned previously. (C) Grid cells develop independent patterns for independent environments. After they are revealed to be part of a single join environment the pattern transform into a coherent global representation [8, 14]. We simulated the patterns independently in two halves of a square to produce incompatible representations (top). After the patterns were spatially stable we continued the learning process of the joined environment (bottom) and both patterns combined into a joined hexagonal grid. (D) For this panel only, we simulate the growth process of a network composed of two subnetworks with a 10% stronger connectivity for cells within each subnetwork (black blocks in the connectivity matrix) compared to the connectivity between subnetworks (yellow blocks). By chance, each subnetwork develops a different solution to the growth process, anchoring to opposite corners of the square enclosure and producing different orientations. Each row shows the average (red) and three individual patterns (green) from each subnetwork. The averages reflect how each subnetwork better represents each corner, with individual patterns showcasing the typical distortions due to the interaction between subnetworks. (E) Stereotypical distortions of the square enclosure [9, 11, 12] include a gradient of orientations (top) and different anchoring solutions (bottom). Experimentally obtained firing maps [11] can be reproduced. Examples are selected by visual inspection from a simulation of 300 grid units stopped at time step 200. (F) Some experimentally reported grid patterns exhibit a long axis that bends backwards [13]. The simulated example (bottom) was selected by visual inspection from a simulation of 300 grid units at timestep 100.

We make an important distinction between two different kind of effects. The first kind are effects solely due to the Turing process and can be also observed in isolated cells not coupled to each other. Effects in this category include rearrangement of field activity (Fig. 4A, top), local lattice defects such as field dislocations (or penta/hepta neighbors) (Fig. 4A, bottom), and the formation of coherent patterns, either by extending an existing one (Fig. 4B, [10, 13]) or merging two stable but incompatible representations (Fig. 4C, [8, 14]). Rearrangement of activity is typical of early times (Fig. 3H), when the initially unstructured distribution of activity transitions into localized fields that slowly position into a triangular lattice. They are usually quickly resolved and are unlikely to last until the end of the learning process. Conversely, lattice dislocations are more likely to become stable, since relocating localized fields into a triangular lattice is much slower than the early clustering of scattered activity into fields. Their origin can be traced back to the uneven spatial distribution and finite population size of their feedforward inputs. However, dislocations are eventually dissolved on a long time scale. Similarly, enough learning time resolves any mismatch between grid patterns that the same cell might have in different areas of the environment, as in the case of merging previously learned environments or revealing novel unexplored areas (Fig. 4B,C). The pattern with stronger feedforward connectivity (stronger anchoring) induces a global restructuring of the pattern with a neighboring field by propagating local interactions starting from the patterns’ overlap and eventually leading to a global coherent reorganization.

The second kind of effects are a direct consequence of the inhibitory coupling. They correspond to geometrical distortions of the patterns and are strongly determined by the boundary of the enclosure. In this group we include general local variations of the pattern’s geometrical properties such as orientation, spacing, and ellipticity (Fig. 4D, [9]), for example the observed gradient of orientations on a single pattern exhibiting shearing (Fig. 4E, top, [11]), different anchoring solutions of the pattern inside a square box (Fig. 4E, bottom), and bending of the grid axis in a highly polarized enclosure such as a trapezoid (Fig. 4F, [12]). Geometrical distortions arise from the competition among patterns that are initially anchored to different locations of the environment. At early times, individual patterns develop with preferred spatial biases due to the initial random distribution of weights and the effect of boundaries, for example different cells may anchor to different corners of a box. These early patterns are slowly transformed into the distorted configurations described above due to interactions via the inhibitory coupling.

Polarized enclosures produce predictable and stereotypical distortions depending on the shape of the boundary. To illustrate this point in the square enclosure, we simulate a network consisting of two equal-size subnetworks whose inhibitory coupling is slightly stronger within subnetworks than between them (Fig. 4D). It allows us to visualize how the interaction between two subnetworks with different grid solutions produces the experimentally observed stereotypical distortions in the patterns (as in Fig. 4E,F). The subnetworks are by chance initially anchored to opposite corners of the square and develop with distinct orientations. The mismatch between the two solutions and the tendency to approach a coherent pattern with time produces the stereotypical distortions of the square enclosure, such as a gradient of orientations [11], different anchoring solutions [12], and a disposition for these distortions to be anchored at opposite corners [9]. In general, uniform random connectivity results in groups of units slightly stronger interconnected than other groups, recreating the biased anchoring of subgroups described.

After reaching triangularity, the other important source of distortions is the organization of the patterns into phase groups. They represent local minima of stability for the growth of individual fields, however, random lateral growth might be strong enough to push individual fields within a pattern towards an adjacent phase group. If a subset of nearby fields within a pattern present a weakly common lateral bias they might transition together producing spatially extended deformations of the patterns. Spontaneous random push and pull of fields and their effect on nearby fields can be observed all throughout the dynamic evolution of the patterns (see supplementary movies). Whenever stabilizing homeostatic factors come into effect, learning stops or slows down resulting in a fraction of cells exhibiting stable geometrical defects and distortions.

From these findings we conclude that the imperfections of experimentally reported grid patterns can be explained by feedforward models with unstructured feedback inhibition.

## III. DISCUSSION

Grid cells form a most efficient code for space [37]. However, geometrical distortions, defects, and environmental influences present in patterns of the local network question the universality of this code [13]. We argued that the existence of these observations is a natural result of the mechanisms leading to triangular patterns in grid cells. In particular, we show that feedforward models are a natural framework to study their development, since they implement a Turing pattern formation process directly over physical two-dimensional space. We found that in these models, during learning, the activation of spatially unstructured feedback inhibition in the grid cell network removes the inherent bias in the weight growth process induced by the presence of physical boundaries, which is well-know for Turing type pattern formation without inhibitory feedback [33]. The model leads to the eventual formation of almost perfect triangular patterns in enclosures of arbitrary shape without any special treatment for the boundaries, such as periodic or fading boundary conditions. Moreover, the model produces a coherent population of grid cells sharing similar orientation, spacing, and field size. In addition, it predicts that grid cells asymptotically self-organize into three phase groups to cover the environment in an approximately uniform way. Finally, the model gives a general account of how interaction among grid cells gives rise to the observed geometrical distortions of the patterns depending on the shape of the recording enclosure and premature termination of learning, and reproduce some of them via unprimed simulations.

Our model assumes that entorhinal cells receive spatially rich inputs. Possible candidates include indirect projections from local layer Vb pyramidal cells, that relay processed information from place cells in area CA1 and subiculum or directly from spatial non-grid cells within superficial layers of mEC [38, 39]. In addition, the model requires cells to be coupled via strong feedback inhibition. In mECII, where most pure grid cells are found, principal cells were reported to be connected to each other exclusively through inhibitory interneurons [38, 40–42]. Stellate cells and pyramidal cells form independent subnetworks that communicate via fast spiking parvalbumin-positive and 5HT3a-receptor expressing interneurons, respectively. Both subnetworks receive direct spatial input from layer Vb pyramidal cells and communicate with each other via intermediary cells. Each subnetwork or their effective combination are good biological candidates to implement the model. Alternatively, the model could as well apply to grid cells in parasubiculum that receive spatially tuned information from thalamic areas or the subiculum [43].

Some feedforward models in the literature already include biological or algorithmic implementations of the principles illustrated here. Kropff and Treves did not employ periodic boundary conditions and yet they could learn triangular patterns in square and circular enclosures [18]. For this, they use additional recovery variables aimed to keep the initially uniform average and variance of patterns constant, which effectively takes the role of an homogeneous inhibitory coupling. It produces an uniform distribution of orientations that is later resolved by the interplay of directional input and additional static but spatially-dependent excitatory collaterals. Stepanyuk offers a mathematical discussion of the effect of non-linearities on the learning process, and in particular how spatially dependent recurrent connections produces a population of equal orientations when input and output are constrained to a twisted torus topology [22].

Continuous attractor network (CAN) models of grid cells also implement a Turing pattern formation process to develop triangular patterns [4]. However, instead of developing the pattern over physical space, the model is implemented on an abstract neural sheet, often with periodic (toroidal) boundary conditions. For conceptual reasons, this model cannot account for effects due to physical boundaries in its simplest form, but might require additional levels of sophistication such as external inputs signaling the location of boundaries [44]. Another important conceptual difference is that all cells in the network reproduce approximately the same unique pattern that is formed in the neural sheet. Depending on the biological implementation, noise can be introduced to weakly alter the patterns from cell to cell [45]. However a rich heterogeneous collection of patterns with defects such as lattice dislocations, splitting of fields, and geometrical deformations are difficult to obtain with a CAN approach. This is in contrast to the observation that only a small fraction of neurons recorded in a local network pass an arbitrary threshold of triangularity, while most of the remaining cells present localized fields of activity and rich spatial information, rather consistent with individual instances of a coupled pattern formation process. CANs are an excellent framework to study relational properties of population responses since they naturally produce populations that share the same orientation, spacing, field size, and coherent phase relationships. However, coherent populations are a signature of strongly coupled systems and do not necessary imply an underlying attractor state, as we have shown using static and spatially unstructured feedback connectivity.

A key feature of the model is the dynamical nature of weights, which change continuously in time. In particular, after reaching triangularity fields may spontaneously drift towards an adjacent phase group. However, in familiar environments grid cell patterns show a high degree of spatially stability on the order of tens of minutes or hours [46]. Although never explicitly modeled, the model assumes that learning slows down or comes to a stop after the animal is familiar with the environment. Additional spatial input such as boundary cells or an over-representation of certain locations by the place cell population might help in anchoring the fields and improving stability. After the initial development period when the patterns reach triangularity, it is not necessarily the case that learning needs to start from scratch when facing a novel environment. For instance, after global remapping almost all place cells change the location of their firing fields, however, the remaining subpopulation maintain a strong bias on certain grid fields that results in a quick rearrangement leading to shifted grid patterns. Since the bias is common to all grid cells, patterns would tend to shift coherently. In general, the speed at which the patterns are learned depends on the particular biological implementation of the model, but in any case a previously established input bias results in a quick rearrangement of grid fields. When facing novel exploration of an environment, an initial input bias helps to anchor a first field of activity in the output cell. The initial field growths stronger with experience and aids in the formation of nearby fields that are later arranged towards a triangular pattern, resulting in fields that seem stable from their first encounter.

A striking property of the class of models discussed in this paper is that grid phases are not random but, at least asymptotically, cluster around three phases yielding an approximately uniform coverage of space. This contradicts reports of homogeneous phase distributions in entorhinal grid cell populations [26]; but see [47]. On the one hand this contradiction can be resolved by assuming that the pattern formation process does not reach its asymptotic equilibrium but learning stops prematurely at a time point at which the three phase clusters are not yet developed (columns 4 or 5 in Fig. 3H). In the same way it could also mean that the spatial input activities *H* are not stationary for long enough time for learning to reach the asymptotic equilibrium. On the other hand the experimental observation that phases in a module are uniformly distributed might be due to undersampling of the local network, and cell pairs used to compute phase relations might not reside in the same locally connected network. An additional caveat is that in the presence of strong distortions and defects the concept of a global grid spatial phase is less useful. An alternative approach is to study for each field in each pattern its spatial relationship to neighbouring fields in all other cells’ patterns, since the model suggests fields from different cells interact only locally in space. Future recordings of modules with a high number of simultaneously recorded cells might be needed to provide a more definite picture, since pooling cells over several sessions recorded over many days might introduce important errors into the phase distribution of patterns.

## IV. MATERIALS AND METHODS

The firing rate approximation

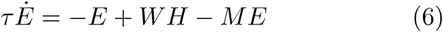

describes the evolution of the cell dynamics. The entries of the inhibitory matrix *M* are i.i.d. random variables drawn from an arbitrary positive distribution with finite variance. We can write it in the form

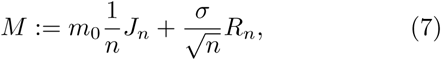

where *J*_*n*_ denotes the square *n× n* matrix whose entries are all equal to one. The matrix *R*_*n*_ is a random matrix whose entries are i.i.d. random variables in the large *n* limit, whose underlying distribution has zero mean and variance equal to one. The scalings were chosen to ensure that the total synaptic strength *m*_0_ and associated variance *σ*^2^ that a cell receives remains unchanged in the large input limit. It additionally provides a fair comparison of the influence of randomness compared to the all-ones matrix for large number of inputs (the circular law ensures the eigenvalues uniformly cover a circle in the complex plane whose radius scales with the variance).

We assume that cell dynamics quickly reach its steadystate compared to the typical times scales of the learning process. The steady-state firing rate

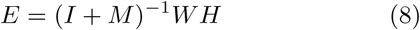

can be rewritten in terms of the mean strength *m*_0_ and the average number of inputs *n* as

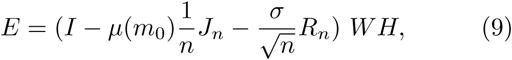

where the 1*/n* scaling in front of the *J*_*n*_ matrix is independent of the particular scaling used in the definition of *M*. The unbiasedness parameter

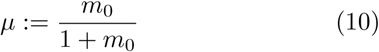

results after taking the approximation of large number of inputs *n*, where we used that the entries of *R*_*n*_ are i.i.d. random variables with zero mean in the large *n* limit.

### A. The learning equation

Without inhibitory self coupling, the weights evolve according to a general Hebbian learning rule supplemented by addittional biological assumptions, which in the large input limit can be written as a continuous approximation

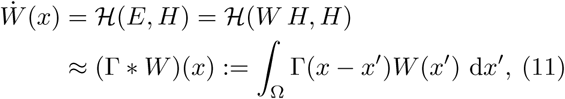

with additional terms needed for convergence of the learning rule to a stable steady-state, typically a nonlinear function depending locally on the weights.

Including self coupling the steady state entorhinal activity *E* is obtained from eq. (9) and thus for a linear Hebbian rule *H*, we have

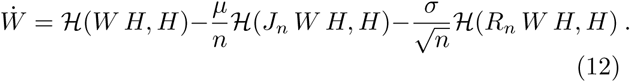

In the large *n* limit this leads to the continuum approximation

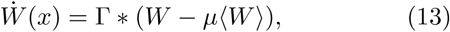

where the angular brackets denote the average over all grid cell patterns *J*_*n*_/*n W* → ⟨ *W*⟩, and 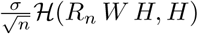 is of order 1 with vanishing expectation value. The scaling of *J*_*n*_ and *R*_*n*_ is chosen such that for fixed *σ* the signal-to-noise ratio is independent of *n*.

### B. The boundary growth bias

The growth bias describes the effect that hard boundaries have on the pattern formation process. It measures how weight changes *δW* (*x′*) in a region Ω affect the growth 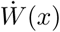 of a weight at a particular location *x*. Formally, the growth bias is thus defined as an integral over the functional derivative of 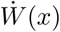

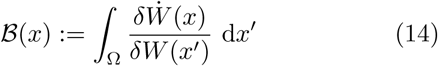

and because of Eq. (3) it amounts to the integral over the shifted kernel,

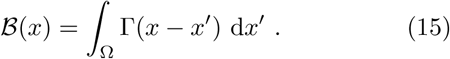

It is strongly determined by the region Ω bounded by the recording enclosure. For a straight infinite boundary, corresponding to the middle parts along a wall of a square recording box, the bias is shown in Fig. 2A as a function of the distance from the wall. For other shapes it was numerically computed and shown in the top row of Fig. 2B.

With the introduction of the inhibitory coupling, the growth bias gets modified as well to

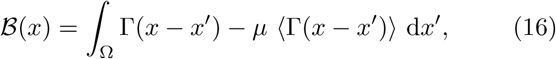

which vanishes for strong inhibition (*µ* ≈ 1), since all grid cells receive the same amount of growth bias, and as a consequence ⟨ Γ = Γ⟩. In general any non-zero value of *µ* weakens the growth bias enough such that triangular patterns are able to develop, albeit in longer times (see Mov. S2,3,5).

### C. Numerical methods

We simulate the development of the weights *W* for about 3000 time steps by the Euler-forward update rule

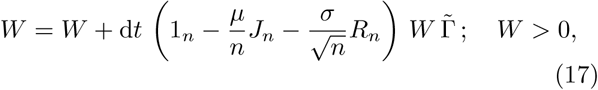

where the parameter values are d*t* = 0.1, *n* = 100, *µ* = 1, and *σ* = 0.2 unless stated otherwise, 1_*n*_ is the identity matrix, and *R*_*n*_ is a Gaussian matrix of zero mean and variance one. W is an *n × m* matrix initialized at random, where *m* is the number of discretization points of the two-dimensional enclosure. It is a square lattice with variable density to ensure the inclusion of approximately 5000 points within the enclosure. For the square this gives the most coarse resolution within the set of used enclosures with a 1.4 cm distance between lattice points (the finest resolution is obtained for the von Neumann elephant, with a 1 cm distance between points). The size of the enclosure is chosen such that the dimension is about one meter, as shown in Fig. 1D. For these points the *m × m* matrix 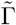 is computed as the value of the learning kernel from Fig. 1B (with maximum plateau value 0.2) for the Euclidean distance between the points. In particular no special treatment is given to points close to the boundary of the enclosure. In all simulations the amplitude of the center region of the learning kernel is always twice that of the surround (never changed), and the surround radius is always twice the radius of the center (*R*_−_ = 2*R*_+_), leaving one degree of freedom in the learning kernel that sets the spacing of the grid. For most simulations *R*_+_ = 5 cm yielding a grid spacing of 16 cm. For Mov. S4,5,7 the center radius is *R*_+_ = 8 cm yielding a grid spacing of 24 cm, and it is varied in Fig. 4 to visually match the experimentally observed grid patterns.

The spacing is obtained from the patterns by computing the distance among all points whose weight value is non-zero and taking the second peak of the smoothed kernel density estimation of the pairwise distances. The spatial phases are obtained as the location of the peak of the field closest to the center of the enclosure. The orientation of a pattern is obtained by computing the orientations of all points in the six neighbouring fields to the center field corrected for the spatial phase. The axes orientations are chosen from the peaks of the circular kernel density estimation from all these orientations. Assuming a perfect hexagonal pattern, the ratio of mean field size to the spacing of a pattern is obtained from the ratio *r* of the number of non-zero weights to the total number of weights as 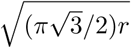.

## ACKNOWLEDGMENTS

The work was funded by the German Research Association (DFG) under grant number LE2250/5-1. M.M. also acknowledges financial support from the Center for Theoretical Neuroscience at Columbia University through the NSF NeuroNex Award DBI-1707398 and The Gatsby Charitable Foundation.

---

Movie S1: Development of patterns for a Spacing of 16 cm. Examples of 5 weight patterns (green) and the average (red) for a population of a 100 patterns. Local fields of activity are formed early on (first hundred time steps) and are first arranged locally in a hexagonal pattern at the population level by anchoring to the top-left corner at around 500 time steps. Local geometrical deformations of the patterns can be observed all throughout the pattern formation dynamics, mostly due to interactions among patterns that induces random lateral growth. Most deformations can be understood as regions of the pattern independently transitioning to a possibly different (locally stable) phase group. Notice the typical defects of the square enclosure reported experimentally, such as different anchoring solutions (Fig. 4E, bottom) in the bottom-middle pattern at around time step 4300, and a gradient of orientations (Fig. 4E, top) in the same pattern at around time step 4600.

Movie S2: Development of patterns for weak connectivity (*µ* = 0.1). Reduced feedback from the population, whose relative strength is measured by *µ*, translates to less random lateral growth in the fields thus greatly improving local spatial stability. Pattern transitions among phase groups leading to geometrical distortions are still possible, for example between timesteps 2000 and 2500.

Movie S3: Development of patterns for weak connectivity (*µ* = 0.5). An intermediate regime of connectivity strength exemplifies the intermediate qualitative behaviour of the patterns in terms of spatial stability and correction of lattice defects.

Movie S4: Development of patterns for a Spacing of 24 cm (*µ* = 1). A simple spatial scaling of the learning kernel Γ results in a corresponding increase for the grid spacing of the patterns. Their development is qualitatively similar to patterns with a smaller spacing such as those in Movie S1.

Movie S5: Development of patterns for a Spacing of 24 cm and weak connectivity (*µ* = 0.1). The qualitative development of patterns is similar to the one shown in Movie S1, with increased spatial stability (compared to Movie S4) due to reduced random lateral growth.

Movie S6: Development of patterns for homogeneous connectivity (*σ* = 0). Noise-free connectivity produces patterns with a uniform distribution of orientations and phases. Patterns present higher spatial stability and therefore are unable (or take longer) to resolve lattice defects developed early on in the pattern formation process, such as a penta/hepta defect in the lower-left corner of the top-left pattern. The probability of random lateral growth in the patterns can be controlled by the parameter *σ* measuring noise in the connectivity.

Movie S7: Development of patterns for a sparse connectivity matrix with only 10% of neurons connected. Patterns break their dependence from the boundary and reach triangularity even for sparsely connected units.

Movie S8: Development of patterns for different enclosures. Patterns reach triangularity independent of the shape of the enclosure. Simple simulations can be used to predict the stereotypical geometric distortions associated to each particular enclosure.

## Notes

### Competing Interest Statement

The authors have declared no competing interest.

## References

[1] T. Hafting, M. Fyhn, S. Molden, M.-B. Moser, and E. I. Moser, Nature 436, 801 (2005).

[2] A. Mathis, A. V. M. Herz, and M. Stemmler, Neural Computation 24, 2280 (2012).

[3] M. Stemmler, A. Mathis, and A. V. M. Herz, Science Advances 1, e1500816 (2015).

[4] M. C. Fuhs and D. S. Touretzky, The Journal of neuroscience 26, 4266 (2006).

[5] N. Burgess, C. Barry, and J. O’Keefe, Hippocampus 17, 801 (2007).

[6] B. Dunn, D. Wennberg, Z. Huang, and Y. Roudi, (2017).

[7] D. Derdikman, J. R. Whitlock, A. Tsao, M. Fyhn, T. Hafting, M.-B. Moser, and E. I. Moser, Nature Neuroscience 12, 1325 (2009).

[8] F. Carpenter, D. Manson, K. Jeffery, N. Burgess, and C. Barry, Current Biology 25, 1176 (2015).

[9] M. Hägglund, M. Mørreaunet, M.-B. Moser, and E. I. Moser, Current Biology 29, 1047 (2019).

[10] F. Savelli, D. Yoganarasimha, and J. J. Knierim, Hippocampus 18, 1270 (2008).

[11] T. Stensola, H. Stensola, M.-B. Moser, and E. I. Moser, Nature 518, 207 (2015).

[12] J. Krupic, M. Bauza, S. Burton, C. Barry, and J. Okeefe, Nature 518, 232 (2015).

[13] J. Krupic, M. Bauza, S. Burton, and J. O’Keefe, Science (New York, N.Y.) 359, 1143 (2018).

[14] T. Wernle, T. Waaga, M. Mørreaunet, A. Treves, M.-B. Moser, and E. I. Moser, Nature Neuroscience, 1 (2017).

[15] M. Gil, M. Ancau, M. I. Schlesiger, A. Neitz, K. Allen, R. J. De Marco, and H. Monyer, Nature Neuroscience 21, 81 (2018).

[16] K. Allen, M. Gil, E. Resnik, O. Toader, P. Seeburg, and H. Monyer, Journal of Neuroscience 34, 6245 (2014).

[17] Y. Burak and I. R. Fiete, PLoS computational biology 5, e1000291 (2009).

[18] E. Kropff and A. Treves, Hippocampus 18, 1256 (2008).

[19] T. D’Albis and R. Kempter, PLOS Computational Biology 13, e1005782 (2017).

[20] L. Castro and P. Aguiar, Biological Cybernetics 108, 133 (2014).

[21] S. N. Weber and H. Sprekeler, eLife 7, e34560 (2018).

[22] A. Stepanyuk, Biologically Inspired Cognitive Architectures 13, 48 (2015).

[23] M. M. Monsalve-Mercado and C. Leibold, Physical Review Letters 119, 38101 (2017).

[24] M. M. Monsalve Mercado, Space in the brain - Of learning and representations, Ph.D. thesis, LMU München: Graduate School of Systemic Neurosciences (GSN) (2018).

[25] A. M. Turing, Philosophical Transactions of the Royal Society B: Biological Sciences 237, 37 (1952).

[26] K. Yoon, M. A. Buice, C. Barry, R. Hayman, N. Burgess, and I. R. Fiete, Nature Neuroscience 16, 1077 (2013).

[27] R. Kempter, W. Gerstner, and J. L. van Hemmen, Physical Review E - Statistical Physics, Plasmas, Fluids, and Related Interdisciplinary Topics 59, 4498 (1999).

[28] W. Gerstner, R. Kempter, J. L. van Hemmen, and H. Wagner, Nature 383, 76 (1996).

[29] C. Leibold, R. Kempter, and L. L. Van Hemmen, Physical Review Letters 87, 248101 (2001).

[30] E. L. Bienenstock, L. N. Cooper, and P. W. Munro, Journal of Neuroscience 2, 32 (1982).

[31] S.-i. Amari, Biological Cybernetics 27, 77 (1977).

[32] K. D. Miller and D. J. C. MacKay, Neural Computation 6, 100 (1994).

[33] J. Horváth, I. Szalai, and P. De Kepper, Science (2009), 10.1126/science.1169973.

[34] T. Solstad, C. N. Boccara, E. Kropff, M.-B. Moser, and E. I. Moser, Science (New York, N.Y.) 322, 1865 (2008).

[35] T. Bjerknes, E. Moser, and M.-B. Moser, Neuron 82, 71 (2014).

[36] J. D. Murray, *Mathematical Biology*, edited by J. D. Murray, Interdisciplinary Applied Mathematics, Vol. 17 (Springer New York, New York, NY, 2004).

[37] A. Mathis, A. V. M. Herz, and M. B. Stemmler, Physical Review Letters 109, 18103 (2012).

[38] M. P. Witter, T. P. Doan, B. Jacobsen, E. S. Nilssen, and S. Ohara, Frontiers in Systems Neuroscience 11, 46 (2017).

[39] G. W. Diehl, O. J. Hon, S. Leutgeb, and J. K. Leutgeb, Neuron 94, 83 (2017).

[40] E. C. Fuchs, A. Neitz, R. Pinna, S. Melzer, A. Caputi, and H. Monyer, Neuron 89, 194 (2016).

[41] E. S. Nilssen, T. P. Doan, M. J. Nigro, S. Ohara, and M. P. Witter, Hippocampus, hipo.23145 (2019).

[42] C. Buetfering, K. Allen, and H. Monyer, Nature Neuroscience 17, 710 (2014).

[43] T. van Groen and J. M. Wyss, Brain Research 518, 227 (1990).

[44] S. A. Ocko, K. Hardcastle, L. M. Giocomo, and S. Ganguli, Proceedings of the National Academy of Sciences of the United States of America 115, E11798 (2018).

[45] O. Shipston-Sharman, L. Solanka, and M. F. Nolan, The Journal of physiology 594, 6547 (2016).

[46] G. W. Diehl, O. J. Hon, S. Leutgeb, and J. K. Leutgeb, Hippocampus 29, 284 (2019).

[47] D. Wennberg, The Distribution of Spatial Phases of Grid Cells, Tech. Rep. (NTNU, Trondheim, 2015).

